# Neural Fingerprinting based on Brain Network Dynamics: A Cross-Platform MEG Study

**DOI:** 10.64898/2026.09.14.751409

**Authors:** Anna Beer, Chetan Gohil, Zoe Tanner, Lukas Rier, Ryan M. Hill, Niall Holmes, Daniel Ferring, Vishal Shah, Mark Woolrich, Matthew J. Brookes

## Abstract

Neural fingerprinting seeks to identify individuals based on measurements of brain activity, exploiting the fact that aspects of brain function unique to an individual remain stable across repeated scans. Magnetoencephalography (MEG) is a powerful technique for fingerprinting. However, most MEG studies have used conventional (SQUID-based) MEG technology and typically rely on data aggregated over time, overlooking the rich temporal dynamics available in MEG. Here, using SQUID-MEG and the more recently introduced OPM-MEG, we showed fingerprinting was possible within and between modalities using static (time-aggregated) features; this is consistent with previous work. We further asked whether fingerprinting was possible based on network dynamics, estimated using a canonical hidden Markov model (CHMM). Results showed that the CHMM-derived networks provide a better fit to SQUID data than to OPM data; however, this effect was small and to be expected given the canonical networks were trained on SQUID data. We further showed that fingerprinting was possible within and between modalities using CHMM-derived state activation time courses and state power spectral densities. However, state summary statistics and transition probabilities only supported within modality fingerprinting. Our findings suggest that subject-specific information is preserved within CHMM states and across MEG technologies. This represents an important step towards advancing our understanding of brain dynamics, and particularly the brain networks delineated by the CHMM. The study also supports the future use of the CHMM for processing and interpretation of OPM-MEG data.

## INTRODUCTION

Traditional neuroimaging focuses on identifying physiological effects that generalise across populations. However, the human brain exhibits substantial inter-individual variability in both its structural and functional organisation, and this has driven a growing interest in understanding differences in physiology between people, how they might relate to behaviour, and their implications for brain health. “Neural fingerprinting” refers to the ability to identify an individual using measurements of brain activity and successful fingerprinting demonstrates that aspects of function unique to that individual are stable across repeated scans (Finn et al. 2015, da Silva Castanheira et al. 2021). As an emerging approach, fingerprinting offers significant potential for developing personalised biomarkers—for example by identifying when disease-related changes disrupt an individual’s characteristic neural signature (da Silva Castanheira et al. 2024). It has also been used as the basis for applications such as brain–computer interfaces, which are often trained based on individuals, not groups (Wittevrongel et al. 2021). Despite the promise, neural fingerprinting remains a new field. Fundamental questions remain regarding which physiological features are most informative for fingerprinting, and how stable these signatures are over time, conditions, and neuroimaging platforms.

Magnetoencephalography (MEG) (Cohen 1972) measures magnetic fields generated by current flow through neural assemblies. Mathematical modelling of these fields enables construction of images showing how brain function changes over time to support cognition (Hamalainen et al. 1993, Baillet 2017). The unique combination of high spatial and temporal resolution afforded by MEG enables not only the ability to localise changes in activity, but also metrics relating to functional connectivity (de Pasquale et al. 2010, Brookes et al. 2011, Brookes et al. 2011, Hipp et al. 2012) and brain networks dynamics (Baker et al. 2014, van Es et al. 2025). This data richness has made MEG a powerful modality for fingerprinting. For example, da Silva Castanheira and colleagues showed that healthy individuals can be identified with high accuracy from just a few seconds of resting-state MEG data (da Silva Castanheira et al. 2021). Supporting this, Wu et al showed that individuals could be identified from MEG data acquired across multiple days, separate tasks, and even between MEG and Electroencephalography (EEG). Clinically, da Silva Castanheira showed that MEG-derived fingerprints are associated with traits of individual patients with Parkinson’s disease (da Silva Castanheira et al. 2024) whilst Sorrentino et al. used connectome-based fingerprinting to show that identifiability is reduced in patients with mild cognitive impairment compared to controls (Sorrentino P et al. 2021).

Most MEG fingerprinting studies have used conventional technology where neuromagnetic fields are assessed using cryogenically cooled sensors based on superconducting quantum interference devices (SQUIDs). However, recent years have seen the adoption of optically pumped magnetometers (OPMs) for MEG (Schwindt et al. 2004, Xia et al. 2006, Schwindt et al. 2007, Shah et al. 2007, Shah and Romalis 2009, Johnson et al. 2010, Johnson et al. 2013, Shah and Wakai 2013, Boto et al. 2017, Brookes et al. 2022, Schofield et al. 2023). OPMs are small, lightweight, and operate without cryogenics. They can be positioned flexibly in arrays tailored to head size (enabling adaptation across the lifespan – babies to adults) (Hill et al. 2019, Rier et al. 2024, Corvilain et al. 2025) and (provided adequate magnetic field control is applied) they can be mounted in helmets that move with the head, enabling data acquisition during movement (Boto et al. 2018, Rea et al. 2022, Sanders BJ et al. 2025). Because OPMs do not require cryogenics they can also get closer to the scalp, generating a higher signal-to-noise ratio (Boto et al. 2016, Iivanainen et al. 2017, Hill et al. 2024) than conventional MEG.

Recent evidence suggests that OPM-MEG is also capable of neural fingerprinting. For example, several papers have performed two or more OPM-MEG recordings in the same individual and showed higher correlation within subjects than between subjects (Boto et al. 2021, Rea et al. 2022, Rier et al. 2023). Rhodes et al. (Rhodes et al. 2023) measured theta-band (4-8 Hz) oscillatory responses to a working memory task and demonstrated successful fingerprinting when contrasting OPM- to SQUID-MEG. Tanner et al. (Tanner et al. 2025) acquired multiple OPM and SQUID datasets in the same individuals and showed that fingerprinting could be carried out both within and between SQUID and OPM platforms. However, these OPM-MEG demonstrations, alongside the majority of SQUID-MEG demonstrations, have employed electrophysiological measurements aggregated over time (for example – spectral power, functional images, connectivity, or time-frequency responses aggregated across all data in a recording). It remains unknown whether the rich, fine-grained dynamics captured by MEG also contain stable, individual-specific signatures that are suitable for neural fingerprinting.

Recent years have seen a step change in our ability to capture brain dynamics using MEG, with the adoption of generative modelling approaches for characterising transient electrophysiological effects. In particular, Hidden Markov Models (HMMs) applied to source-reconstructed MEG data enable the identification of recurring spatiotemporal patterns of electrophysiological activity (Baker et al. 2014, Quinn AJ et al. 2018, Vidaurre D et al. 2018, Vidaurre D et al. 2018). These patterns, typically referred to as *brain states*, represent distributed functional networks, each characterised by a distinct spatial signature, functional connectivity profile, and spectral content. HMMs characterise dynamics on a millisecond timescale. They provide the probability of being in a particular brain state at each time point as well as the probability of transitioning between states.

Repeated application of HMMs across resting-state and task-based MEG studies has demonstrated that many of these transient states are highly reproducible, leading to the development of a canonical Hidden Markov Model (CHMM) (Gohil C et al. 2026). Using this approach, rather than fitting a new HMM to each individual dataset, the CHMM provides a common repertoire of brain states, derived from thousands of MEG recordings, that can be applied to new datasets, enabling consistent characterisation of brain network dynamics across studies. However, an open question is whether this CHMM is transferable to OPM-MEG data, or whether the rapid state dynamics that it uncovers can be used for neural fingerprinting.

Here, we analyse SQUID- and OPM-MEG data from a previous study (Tanner et al. 2025) to address three questions. First, can we replicate the finding that fingerprinting is possible using conventional *static* (time-averaged) data features, both within and between MEG platforms (Tanner et al. 2025)? Second, can a network-dynamics-based approach (CHMM) trained exclusively on SQUID-MEG data generalise to OPM-MEG data, with a comparable quality of fit? Finally, can features describing these dynamic networks be used to achieve neural fingerprinting, both within and across MEG platforms?

## METHODS

### Participants

The data used in this study have been reported previously (Tanner et al. 2025). In total there were 60 datasets recorded from 15 participants (7 male, 8 female; mean age 27 ± 5 years; 14 right-handed). Each participant undertook 4 scans, 2 using OPM-MEG and 2 using SQUID-MEG. All subjects had an anatomical MRI scan, which was used for source modelling (see below). Data were acquired at the Sir Peter Mansfield Imaging Centre, University of Nottingham, UK. All participants gave written informed consent prior to taking part, and the study was approved by the University of Nottingham Faculty of Medicine and Health Sciences Research Ethics Committee (approval number H16122016).

### MEG systems

The OPM-MEG system comprised 64 triaxial sensors (QuSpin, Colorado, USA) each measuring magnetic field along 3 axes (Boto et al. 2022) – meaning the system comprised 192 independent channels. Sensors were mounted in a 3D printed helmet (Cerca Magnetics Limited, Nottingham, UK) and controlled by a miniaturised integrated electronic control system (Schofield et al. 2024) (*“NEURO-1”*, QuSpin, Colorado, USA). The system was housed in a 5 layer magnetically shielded room (MSR) (Magnetic Shields Limited, Kent, UK). (4 layers of MuMetal help reduce low frequency magnetic interference and 1 layer of copper helps reduce high frequency interference.) The MSR was also equipped with degaussing coils to demagnetise the MuMetal layers (Altarev et al. 2014). A matrix coil system (Holmes et al. 2023) (Cerca Magnetics Limited, Nottingham, UK) was used to control background field inside the MSR.

The SQUID-MEG comprised 275 radially oriented axial gradiometer pick up coils (5 cm baseline) coupled to 275 SQUIDs (CTF, Vancouver, BC, Canada). The system also incorporated a reference array with a further 29 SQUID-based sensors located distal to the head. This enables rejection of background interference via the construction of 3^rd^ order synthetic gradiometers (Vrba and Robinson 2001). The system was housed inside an MSR comprising 2 layers of MuMetal and 1 layer of aluminium (Ak3b, Vacuumschmelze, Hanau, Germany).

### Experimental paradigm and data collection

The task involved sensory stimulation of the left and right index fingers using 2 “braille” stimulators. Each stimulator contained 8 “pins” arranged in a 4 x 2 matrix. Each pin could be positioned “up” (stimulating) or “down” (not stimulating) and by manipulating which pins were up, “braille-like” tactile patterns could be applied to the participants’ fingers. Here two patterns were used: a *non-target* in which all eight pins were up, and a *target* in which only the top two pins were up.

A single trial began (at *t = 0*) with an auditory attentional cue telling participants to either “attend left” or “attend right.” Following this, at *t=1.17 s* a series of 5 braille patterns were presented, at intervals of 1.37 s, to both index fingers. The stimuli lasted 370 ms with a 1 s pause between patterns. Participants were asked to respond, with a button press using both their left and right thumb, if the target pattern was presented to the attended hand. Each trial ended with a 7-s rest period. A single experiment comprised 80 trials. The probability of a target was 0.2.

The paradigm was identical in both MEG systems, and each participant was scanned 4 times in a single day; twice in the morning and twice in the afternoon, with a ∼2-hour break in between the morning and afternoon sessions. The order of the scanning sessions was counterbalanced with 8 participants having OPM-MEG in the morning and SQUID-MEG in the afternoon, and the other 8 having SQUID-MEG in the morning and OPM-MEG in the afternoon. In each session, the participant was sat comfortably with their head in the MEG helmet whilst data were recorded throughout the paradigm (with a sampling rate of 375 Hz for OPM-MEG and 600 Hz for SQUID-MEG). For OPM-MEG, at the start of each session the room was demagnetised and the matrix coil used to reduce background field to <1nT.

### Coregistration of MEG data to brain anatomy

For OPM-MEG, immediately following data acquisition, a structured light camera (Einscan H, SHINING 3D, Hangzhou, China) was used to gather 3D images of the participant’s head and face as well as the helmet (in situ). The 3D surfaces representing the head and face were extracted from these scans and fitted to equivalent surfaces extracted from the participant’s anatomical MRI scan. This, coupled with knowledge of the locations of sensors in the helmet (from the 3D printing process) enabled coregistration of the sensor geometry to the participants brain anatomy.

For SQUID-MEG, 3 head position indicator (HPI) coils were attached to the participant’s head; these were energised during the scan and a magnetic dipole fit used to determine their locations relative to the MEG sensors. Following data acquisition, the locations of the HPI coils were digitised relative to the subject’s head shape using a 3D digitiser (Polhemus, Vermont, USA). The head/face surface from this digitisation was fitted to the equivalent surface extracted from the MRI scan, and this, coupled with knowledge of sensor locations relative to the HPI coils, enabled coregistration.

### Data preprocessing

Data from each session were pre-processed using the osl-dynamics (osld) toolbox (Gohil C et al. 2024), implemented in Python. OPM-MEG and SQUID-MEG data were treated equivalently (unless otherwise stated below).

First, all data were down-sampled to 250Hz and bandpass filtered between 1-120Hz using a 5th order Butterworth filter. Notch filters at 50 and 100Hz were applied to remove artefacts generated by mains electricity. (For the SQUID-MEG datasets, additional spikes at 57Hz and 109 were also removed using a notch filter.) Data were then segmented into trials; trials began at t = 0s (at the onset of the auditory attentional cue) and lasted 13.65s.

Power spectral density plots were created for all channels, and those exhibiting significantly elevated variance were identified using the generalised extreme studentised deviate (G-ESD) test. Guided by these detections, the data identified were inspected visually and channels exhibiting excessive noise, or abnormally low variance indicative of sensor malfunction, were removed.

Time-series data for each session were also screened for segments containing artifacts using the G-ESD test. Marked segments containing artifacts were subsequently reviewed by visual inspection and removed if necessary. All subsequent analyses employed all remaining data after bad segment removal; however, for any analysis requiring trial averaging, trials with bad segments were removed in their entirety.

Finally, a further 4–45 Hz band-pass filter was applied to the cleaned data using a 5^th^ order Butterworth filter.

### Source Reconstruction and Parcellation

To enable source reconstruction, the brain space was divided into a regular 8 mm grid of voxels, and an estimate of the electrophysiological activity was made at each voxel using a linearly constrained minimum variance (LCMV) beamformer (Robinson and Vrba 1998). The beamformer creates a source space estimate of neural current using a weighted sum of magnetic field measurements made at each sensor. The weights are derived to minimise the variance of the output signal, subject to a linear constraint that variance originating at the location of interest (i.e. a specific voxel) must be maintained. This requires a model of the magnetic fields that would be generated by a current source at each voxel (i.e. the forward model), which was computed based on a single-layer volume conductor, and a current dipole model (Nolte 2003). Weights calculation also requires computation of data covariance, which was based on the filtered data from all trials (excluding bad channels and segments). The data covariance matrix was regularised using the Tikhonov method with a regularisation parameter equal to 5% of the maximum eigenvalue of the unregularized matrix. For each voxel, the source orientation was taken as that which maximised projected power. This resulted in a single time course of electrophysiological activity per voxel. Reconstructed source estimates were subsequently transformed into MNI space.

Voxels were appointed to anatomical parcels according to the Glasser parcellation (Glasser et al. 2016) which defines 52 anatomical regions. Within each parcel, voxel-wise source estimates were aggregated with the aim of deriving a single representative electrophysiological timecourse per parcel. Specifically, timecourses from all voxels within a parcel were concatenated in the voxel dimension, and the first principal component (PC) taken. The PC timecourses for each parcel were then rescaled to ensure that their standard deviation was equal to the weighted average of the standard deviations of the voxel timecourses in that parcel. The resulting scaled PC was taken as the parcel timecourse.

*“Static” representations of neural activity*

To derive a “static” (i.e. averaged over time) metric of neural activity for each parcel, we calculated the power spectral density (PSD) of the parcel time courses (in the 4–45 Hz frequency range). PSDs were calculated independently for each parcel and session, allowing the PSD to be compared across regions, participants, sessions and recording modalities.

### Canonical Hidden Markov Model

The HMM has been explained in detail in previous publications (Baker et al. 2014, Quinn AJ et al. 2018, Vidaurre D et al. 2018, Vidaurre D et al. 2018). Briefly, the variant used here is based on a “time-delay embedded” (TDE) HMM, developed to identify transient patterns of oscillatory neural activity from parcellated MEG data. The TDE-HMM appends extra time-lagged copies of the original parcellated MEG time courses to form a higher-dimensional time series. These extra channels enable each state’s covariance matrix to capture not only patterns of regional variance/covariance but also the temporal auto/cross-correlation between embeddings. This allows identification of frequency-specific networks. The model infers a set of recurring brain *states*, each characterised by a spatial distribution of oscillatory power, PSD and functional connectivity. Additionally, the HMM provides a probabilistic time course describing when each state is active, and a transition probability matrix that characterises the probabilities of transitioning between each state.

In the canonical Hidden Markov Model (CHMM) (Gohil C et al. 2026), a time-delay embedded HMM has been pre-trained on a large normative dataset of healthy participants, recorded using SQUID-MEG (specifically, the “CamCAN” dataset (Taylor JR et al. 2017)) during both resting-state and task-based experiments. The CHMM defines eight states (specifying the covariance matrix of each state and a state transition probability matrix). These states capture recurring patterns of large-scale electrophysiological activity observed across a population. When analysing a new dataset, rather than inferring a completely new set of states (as with a traditional HMM) the CHMM takes the canonical states from the normative dataset and estimates the probability that each canonical state is active, at every time point in the new dataset. The inferred probability time courses are then converted to a binary time course by identifying the state with the maximum probability, at each point in time.

To apply the CHMM, our parcel data were orthogonalised to remove zero-phase lag correlations between parcel time courses (to conservatively remove source/spatial leakage) (Colclough et al. 2015). The data were then time delay embedded, reduced in dimension via a principal component analysis, and z-scored before the CHMM was inferred. The following measures were then derived:

- **Variational free energy (VFE)** quantifies how well the CHMM model explains the observed data while accounting for model complexity. In practice, VFE is commonly used as a goodness-of-fit metric with lower free energy values indicating that the inferred state structure captures the data more effectively.
- **State power maps.** The timecourse of signal power for each parcel is derived, and then weighted according to the probability of each state. These data are then summed over time. The result is 8 maps of power, specific for each state. These are then normalised by the mean power for each parcel, computed over all time and states. The final result is a map of relative state power: For any given region and state, a positive value means that region has a higher than average power during state occurrence; a negative value means a lower power.
- **State PSDs.** State-specific PSDs are estimated by weighting the original parcel time series according to the inferred state probabilities. Time points with a higher probability of belonging to a given state then contribute more strongly to that state’s spectral estimate. Parcel-wise PSDs are then averaged to obtain a whole-brain PSD for each state.
- **Trial-averaged state activation time courses.** The binary state activation sequences are summed over trials, and the result is divided by the number of trials. This results in trial time courses showing the probability of each state being active as a function of time.
- **Summary statistics of the state dynamics.** Summary statistics are calculated from the binary state time courses and include:

1. **Fractional occupancy** - the proportion of the total time spent in each state
2. **Mean lifetime** - the average duration of a continuous state visit
3. **Mean interval time** - the average duration between successive visits to a state
4. **Switching rate** - the average number of state activations per second
- **State transition probabilities.** Empirical transition probability matrices are calculated from the inferred binary state sequence by counting transitions between specific pairs of states and normalising by the total number of transitions originating from each state. These empirical transition matrices vary between recordings and can therefore be used to characterise individual differences in state-switching behaviour.

### Comparing CHMM fit across scanner modalities

To assess whether the CHMM model (which was trained on SQUID data) generalised to our SQUID-derived data (from a scanner with a different architecture to the training data) and our OPM-MEG data, we used the VFE. Specifically, VFE was calculated for each of our SQUID and OPM datasets, and this was compared to the VFE values derived from the original training data. To determine whether the mean VFE of the SQUID and OPM data differed significantly, a permutation test was performed: 30 datasets from the original CHMM training set were selected randomly, and their mean VFE was calculated. This process was repeated 1,000,000 times to generate a null distribution. The mean VFE values from our SQUID and OPM data were then compared against this distribution.

### Neural fingerprinting

The principle of neural fingerprinting is that within-subject correlation of brain activity should be higher than between-subject correlation. To quantify this, all 60 datasets were processed using the static (PSD) and CHMM analyses and the following 5 fingerprinting metrics were extracted:

- **Static PSD correlation:** The PSD for each parcel in the first dataset was correlated with the corresponding parcel PSD in the second dataset. This produced 52 correlation coefficients, which were averaged over parcels.
- **State timecourse correlation:** The trial averaged state activation timecourses for each state in the first dataset was correlated with the corresponding timecourse in the second dataset. This resulted in 8 correlation coefficients (one per state) which were averaged.
- **Summary statistic correlation:** The values of each statistic (fractional occupancy, mean lifetime, etc) for each state were concatenated to give 4 vectors, each of dimensions 1 x 8. These vectors were correlated between datasets, and the resulting 4 correlation coefficients averaged.
- **State PSD correlation**: The PSD associated with each state was correlated between datasets and the resulting eight correlation coefficients averaged.
- **State transition correlation:** The 8 state transition probability vectors (i.e. each showing the likelihood that a specific state transitions into every other state) were correlated between datasets (with self-transitions excluded). The resulting 8 correlation coefficients were averaged.

The above quantities were computed for every possible pair of datasets, resulting in a set of 5 metrics per pair.

To investigate neural fingerprinting, we aimed to construct a set of fingerprinting matrices where the on-diagonal elements represent within subject correlations, and the off-diagonal elements represent between subject correlations. However, this study is complex since we had four recordings per participant (two OPM-MEG sessions and two SQUID-MEG sessions). We therefore followed a previous approach (Tanner et al. 2025). To explain, assume we have 2 subjects (labelled M and N) and 4 sessions per subject (labelled OPM1, OPM2, SQUID1 and SQUID2).

### Within-platform fingerprinting

There are two possible within-subject comparisons per modality (e.g. M_OPM1_ to M_OPM2_ and N_OPM1_ to N_OPM2_). However, there are four possible between-subject within-modality comparisons (e.g. M_OPM1_ to N_OPM1_; M_OPM1_ to N_OPM2_; M_OPM2_ to N_OPM1_ and M_OPM2_ to N_OPM2_); though only two of these are independent. To ensure fairness, we therefore chose to only keep two of the four between subject comparisons (M_OPM1_ to N_OPM2_ and M_OPM2_ to N_OPM1_). This meant that we could construct within-modality fingerprinting matrices where each matrix element represents a single comparison between 2 datasets.

### Cross-platform fingerprinting

For cross-platform fingerprinting, there are always four possible within-subject comparisons (e.g. M_OPM1_ to M_SQUID1_; M_OPM1_ to M_SQUID2_; M_OPM2_ to M_SQUID1_ and M_OPM2_ to M_SQUID2_;). Similarly, there are four between-subject comparisons (e.g. M_OPM1_ to N_SQUID1_; M_OPM1_ to N_SQUID2_; M_OPM2_ to N_SQUID1_ and M_OPM2_ to N_SQUID2_;). Since in all cases the number of comparisons is the same, we averaged all four values together to construct each element of the cross-platform fingerprinting matrices. More detail on this approach can be found in Tanner et al 2025 (Tanner et al. 2025). Fingerprinting matrices were constructed, within and across platforms, for each of the 5 correlation measures listed above.

Having derived the subject-by-subject correlation matrices, we conducted two analyses. First, we assessed how many of the 15 subjects could be successfully identified for each metric (i.e. for each row in the fingerprinting matrix; how many times does the maximum correlation value fall on the leading diagonal). Second, to quantify the overall separation between within-subject and between-subject correlations, we calculated the difference between the mean within-subject correlation (mean diagonal value) and the mean between-subject correlation (mean off-diagonal value). This measure asks: how much more similar are subjects to themselves, than to the average of all other subjects? Positive values indicate that, on average, within-subject correlations exceed between-subject correlations. We term this metric the *average correlation margin*. It is worth noting that this metric quantifies the extent to which the correlated measure contains subject-specific information, rather than the extent to which individual subjects can be uniquely identified.

To assess statistical significance of the average correlation margin, we used a permutation test. We reasoned that if no subject-specific information existed in the MEG metrics, then the order of the elements in the fingerprinting matrices could be randomised without changing the average correlation margin. We therefore randomised the structure of the correlation coefficient matrices and rederived the average correlation margin. This process was repeated 1,000,000 times to create a null distribution. Then, by comparing the real average correlation margin (without randomising the correlation matrix) to the null distribution we derived an empirical p-value. The threshold for significance was set at 0.05, however this was corrected for multiple comparisons (using the Bonferroni technique). Specifically, we tested fingerprinting using 5 metrics (static PSDs; state timecourses; state PSDs; summary statistics and state transition); for each metric we derived an average correlation margin in 3 ways (OPM-OPM; SQUID-SQUID and OPM-SQUID). We therefore Bonferroni-corrected for multiple comparisons across 15 tests (i.e. α = 0.0033).

## RESULTS

Figure 1a shows group-level static power maps depicting the spatial distribution of theta (4-8Hz), alpha (8-13Hz) and beta (13-30Hz) power. In all cases, the mean power level (across all parcels) has been subtracted (hence the negative values) and data for OPMs (top) and SQUIDs (bottom) are shown. Data are averaged across participants and experimental runs within each modality. The spatial distribution of power across parcels is as expected; for example, the largest alpha power is in the occipital regions. There is also a compelling similarity between OPMs and SQUIDs. The right-hand side shows representative parcel PSDs; the case on the left shows the occipital lobe and the case on the right shows the frontal lobe. In both cases, data are averaged across datasets for each modality. Note a clear spectral difference between regions, but strong similarity between modalities.

**Figure 1:**
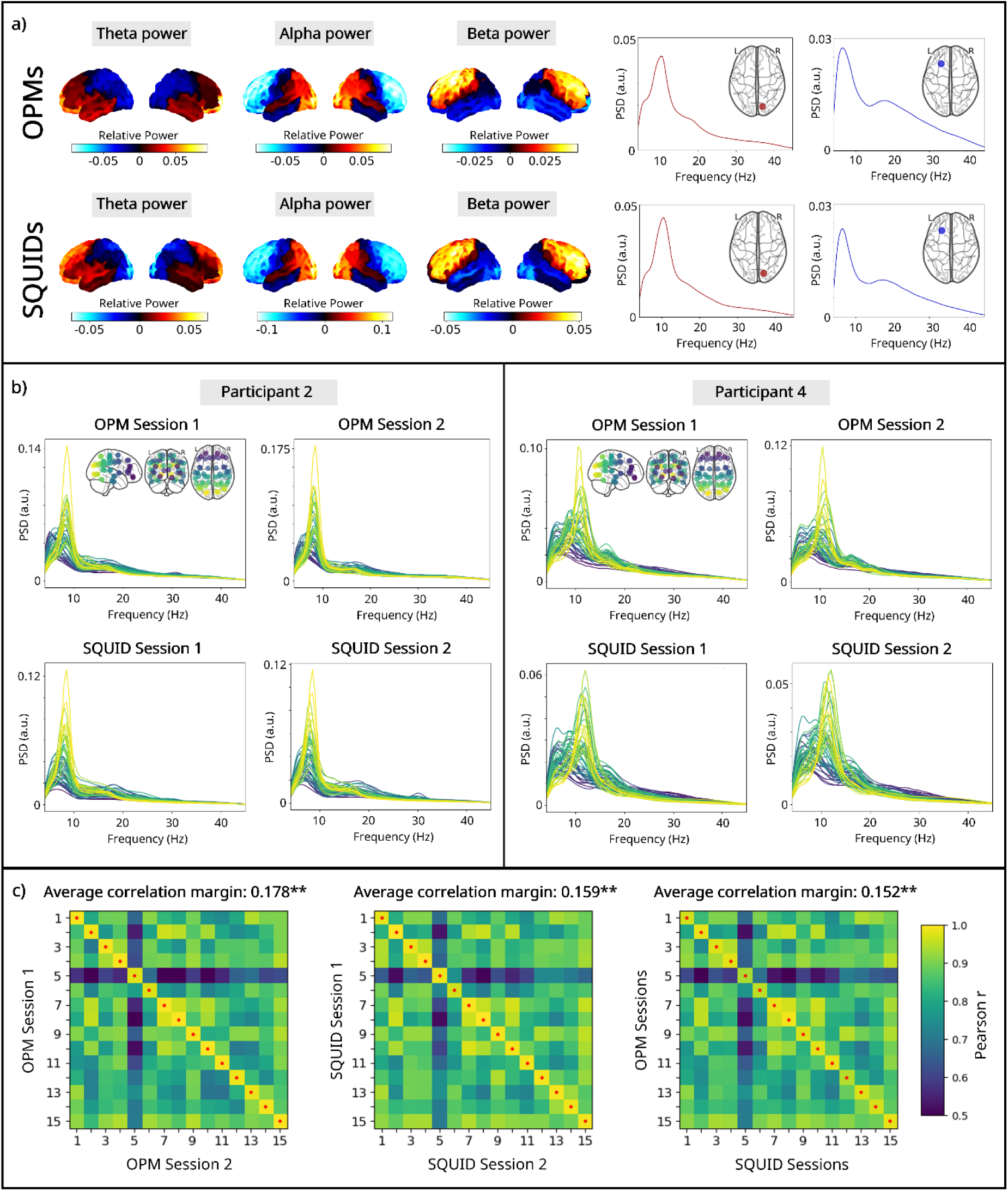
**Neural fingerprinting using static data**. a) left: Group-average source power maps for the theta (4–8 Hz), alpha (8–13 Hz), and beta (13–30 Hz) frequency bands. Note that the mean power across all brain regions has been subtracted. Right: example PSDs from regions in frontal and occipital lobes; data averaged across subjects. b) Source-level PSDs from all 52 parcels for two representative participants. PSDs from all parcels are overlaid and coded by colour (see inset plots). c) Fingerprinting matrices comparing parcel PSDs within OPM recordings (left), within SQUID recordings (middle), and across OPM and SQUID recordings (right). Red dots show the highest correlation value per row. Note highest correlations are within subjects (i.e. fall on the diagonal). Average correlation margin values are shown above each matrix; ** indicates significance following correction for multiple comparisons using Bonferroni correction.

Figure 1b shows example regional PSDs (all parcels overlaid) for two participants (2 and 4). Data are shown for all four recording sessions, and the colours relate to different parcels (key inset). Note that the spectral profiles are consistent across both recording sessions within a modality and between modalities. However, there is a marked difference between participants.

Figure 1c shows fingerprinting matrices based on static PSDs. The left panel shows OPM-to-OPM fingerprinting; the middle panel shows SQUID-to-SQUID, and the right panel shows OPM-to-SQUID. In all cases, matrix element [*i*, *j*] represents the correlation between participant *i* and participant *j*; diagonal elements correspond to within-subject correlations. For all three matrices and for all 15 subjects, the highest correlation falls on the diagonal (indicated by the red dot), demonstrating that each subject was most strongly correlated with themselves, and could therefore be correctly identified. The average correlation margins were highest for the OPM-OPM comparison and lowest for the between modality comparison (as might be expected). However, all three values were significant according to a permutation test. We also see similarity of the matrix structure across the OPM-OPM, SQUID-SQUID, and OPM-SQUID matrices. These data agree with previous findings (Tanner et al. 2025) in showing that static measures enable successful fingerprinting.

Figure 2 shows histograms of VFE values for the training datasets compared with equivalent histograms for our SQUID (panel a) and OPM (panel b) data. The mean VFE was 166.2 for the training data, 166.2 for our SQUID data, and 167.2 for our OPM data. The mean VFE of the OPM MEG data was significantly (p < 0.05 based on a permutation test) higher than that of both the training data and our SQUID data. However, the VFE values themselves for the OPM data still fell within the range that was observed for the training data. This important finding will be addressed further in our discussion.

**Figure 2:**
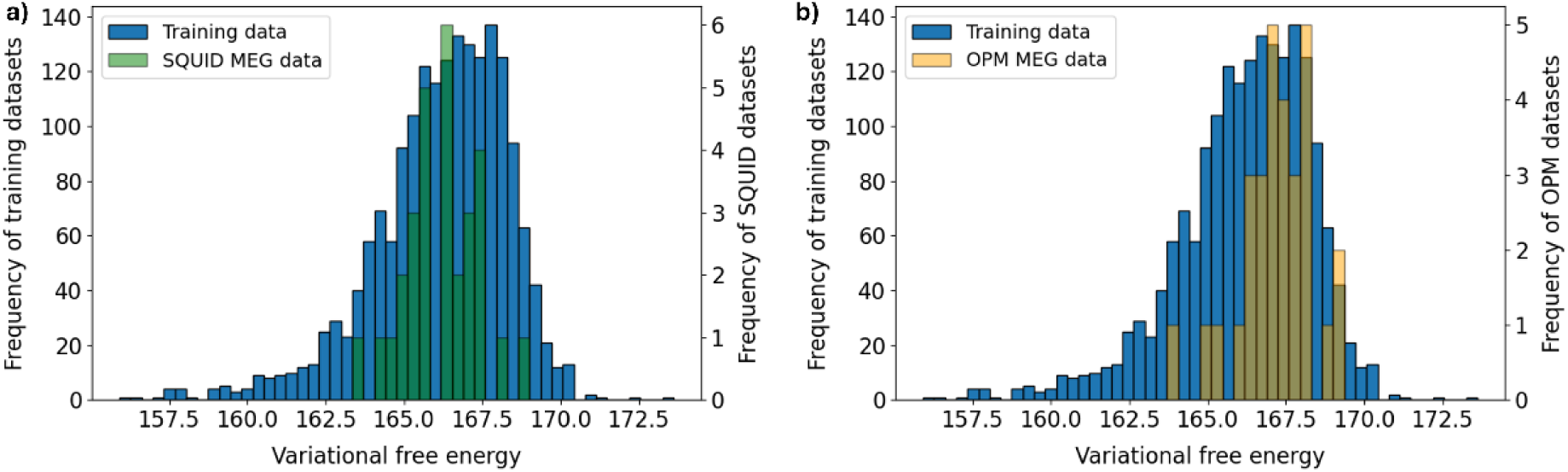
VFE for SQUID and OPM data. a) Histogram of VFEs for the training data (blue) with SQUID data (green) overlaid. b) Histogram of VFEs for the training data and OPM data.

Figure 3 shows the CHMM state power maps (left) and trial-averaged state activation timecourses (right) for the eight canonical states. Data from OPM and SQUID recordings are shown, averaged over participants and experimental runs within each modality. Recall that the power maps are shown relative to overall power (hence contain negative values). The dashed vertical lines in the timecourses indicate the onset of the five Braille stimuli.

**Figure 3:**
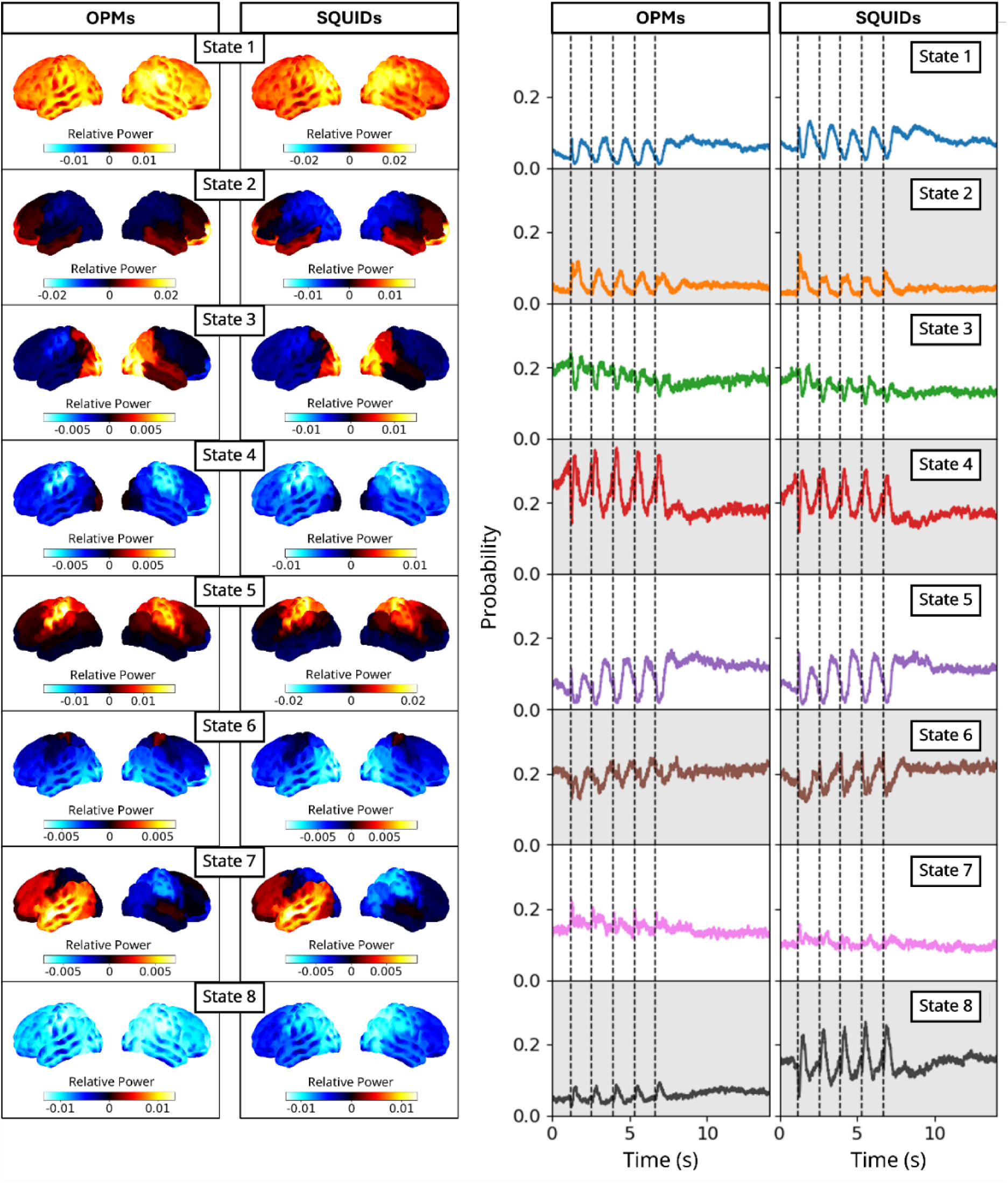
CHMM applied to OPM and SQUID data. Left: Power maps for the eight canonical states inferred from our two recording modalities. All maps are averaged across 15 participants and both experimental runs per participant (for the same modality). Left hand plots show OPMs, right hand plots show SQUIDs. Right: Trial-averaged state activation timecourses, averaged over both runs and participants. Note these are absolute values of probability (i.e. timecourses are shown with no baseline correction). Vertical dashed lines indicate the onset of the five Braille stimuli.

The spatial signatures of the power maps are in good agreement across modalities, however, recall that this is to be expected because the states are imposed by the CHMM. The timecourses show that all states modulate with the task, with the most pronounced modulation in states 5 and 6, which are linked to the sensorimotor cortices – as expected given the task. There is good correspondence between OPM and SQUID derived timecourses (which is not imposed by the model); though there are also differences (e.g. the higher probability of occurrence of state 8), which will be discussed below.

Figure 4a shows state-specific PSDs for the 8 inferred states, for two representative participants. Data are shown for all four sessions. It is noteworthy that each state exhibits a distinct spectral profile that is preserved across repeated OPM and SQUID recordings. However, there are noticeable differences between participants suggesting that, like the parcel PSDs shown in Figure 1, the inferred state spectra retain characteristics relating to a single individual. Figure 4b shows the corresponding fingerprinting matrices. Again, we see strong diagonal structure for all three matrices. In the OPM-OPM (left) and SQUID-SQUID (middle) cases, all participants are correctly identified. For the OPM-SQUID case, 13 out of 15 participants could be correctly identified. The average correlation margin was similar for all 3 matrices and all three average correlation margin values were significant according to the permutation test, suggesting that the state spectra retain individual characteristics, even when measured using different modalities. (As an aside, note that whilst it might be tempting to compare average correlation margin scores between this metric and that presented in Figure 1 (static PSD correlation), this would be invalid since the correlation values (hence the average correlation margin values) derived are based on a different number of degrees of freedom.)

**Figure 4:**
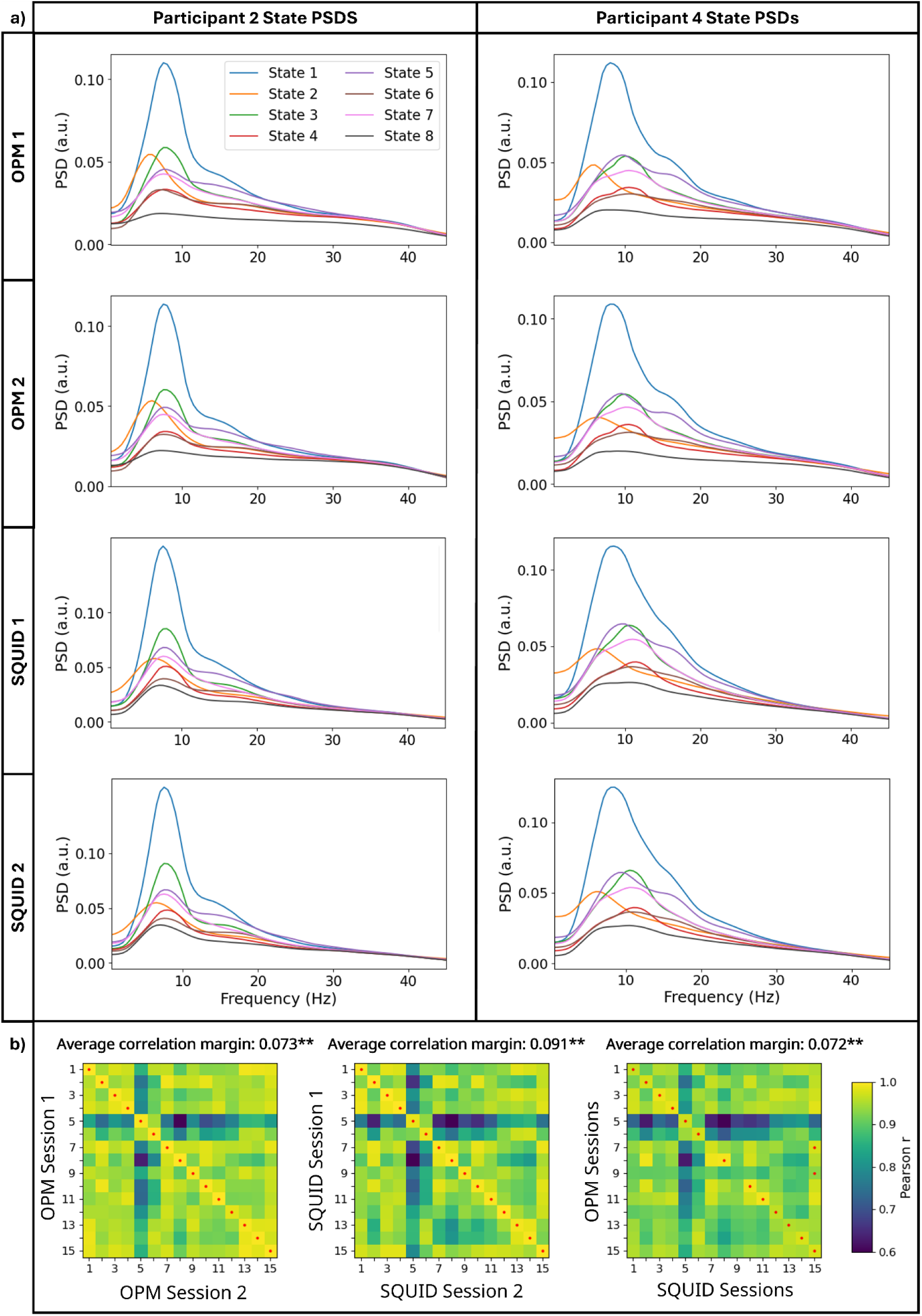
Fingerprinting using CHMM canonical state spectra: a) State PSDs for the eight inferred states from two representative participants. Data for two OPM and two SQUID recording sessions shown. b) Fingerprinting matrices comparing state PSDs within OPM recordings (left), OPM recordings (left), SQUID recordings (middle), and between modality recordings (right). Red dots again show the highest correlation value per row. Average correlation margin values are shown above each matrix; ** indicates significance following correction for multiple comparisons using Bonferroni correction.

Figure 5a shows the trial-averaged state activation probability time courses for two representative subjects, across the four recording sessions. For each subject, the temporal profiles of all 8 inferred states are shown. Once again, these are relatively consistent across repeated recordings and between the OPM and SQUID systems. However, there is a high between-participant difference. For example, in state 6, participant 2 consistently shows that the probability of state occurrence drops during stimulation and then increases on stimulus cessation. However, participant 4 consistently shows almost no change in state probability throughout the trial.

**Figure 5:**
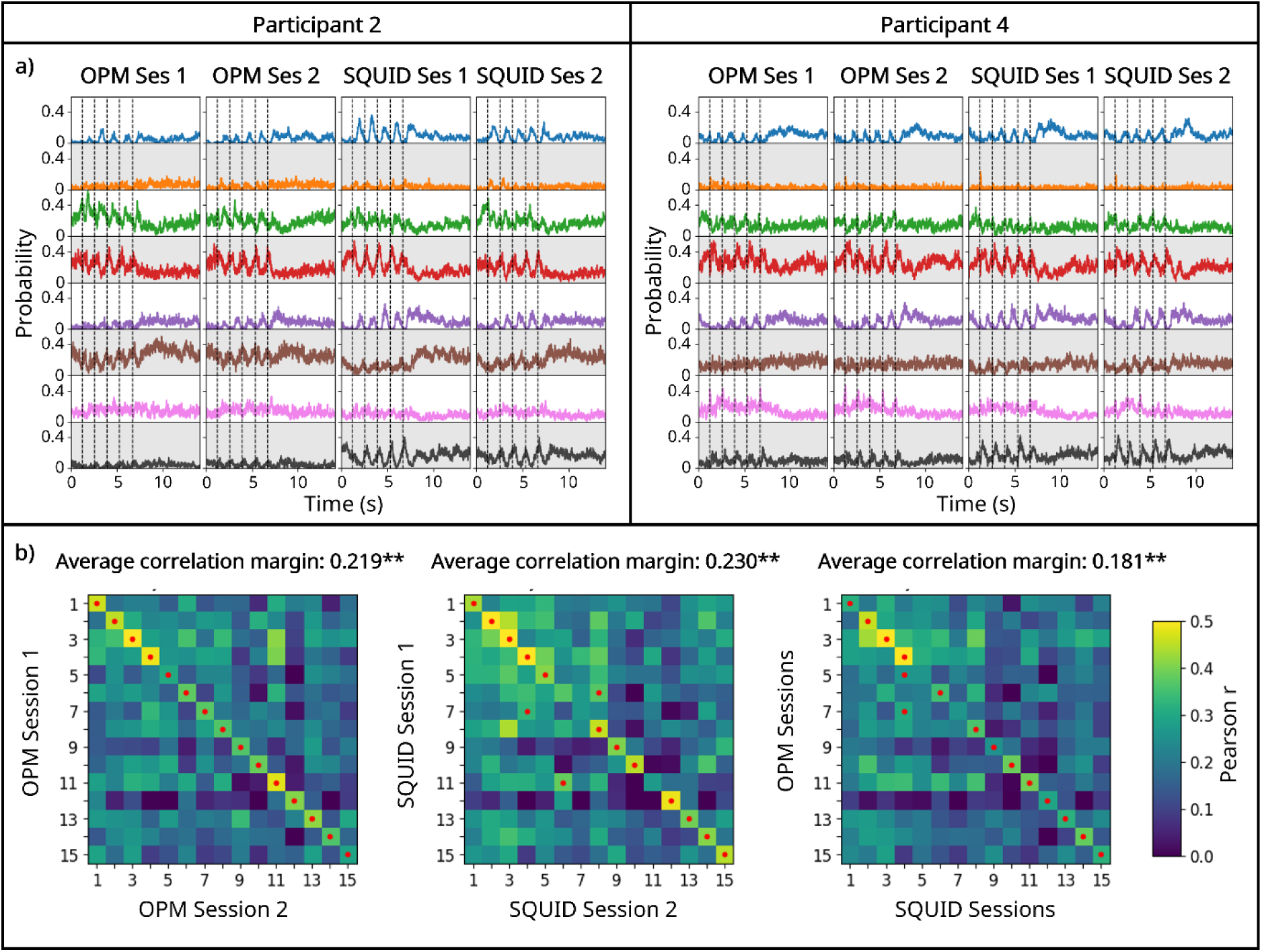
Fingerprinting using CHMM trial-averaged state activation timecourses: a) Trial-averaged state activation timecourses (i.e. stimulus-locked state probability time courses) for the eight inferred states, in two representative participants. Data from all four recording sessions shown. The vertical dashed lines indicate Braille stimulus onset. b) Fingerprinting matrices comparing state activation time series; OPM recordings (left), SQUID recordings (middle), and between modality recordings (right). Average correlation margin values are shown above each matrix; ** indicates significance following correction for multiple comparisons using Bonferroni correction.

The fingerprinting matrices shown in Figure 5b support this observation, showing a clear diagonal structure for the OPM-OPM, SQUID-SQUID, and OPM-SQUID comparisons. Here, all 15 subjects were identified correctly for the OPM-OPM comparison; 12/15 were correctly identified using SQUIDs, and 13/15 were identified correctly for the OPM-SQUID comparison. We see significant average correlation margins for all three comparisons, with the highest for SQUID-SQUID and the weakest for between modality comparison. (Although this may appear inconsistent with the identification accuracies, the average correlation margin reflects the average separation between within-subject and between-subject correlations rather than individual identifiability. Consequently, a higher margin can arise from lower average between-subject similarity even when identification accuracy is unchanged or lower.) Together, these findings demonstrate that, like the state-specific spectra, the state probability time courses contain participant-specific information that is preserved across modalities.

Figure 6 shows the standardised (z-scored) state summary statistics for the inferred states. Figure 7 shows the empirical state transition probability matrices derived from the binary state timecourses. In both Figures, panel a shows representative examples for the two representative participants, whilst panel b shows the corresponding fingerprinting matrices for all subjects.

**Figure 6:**
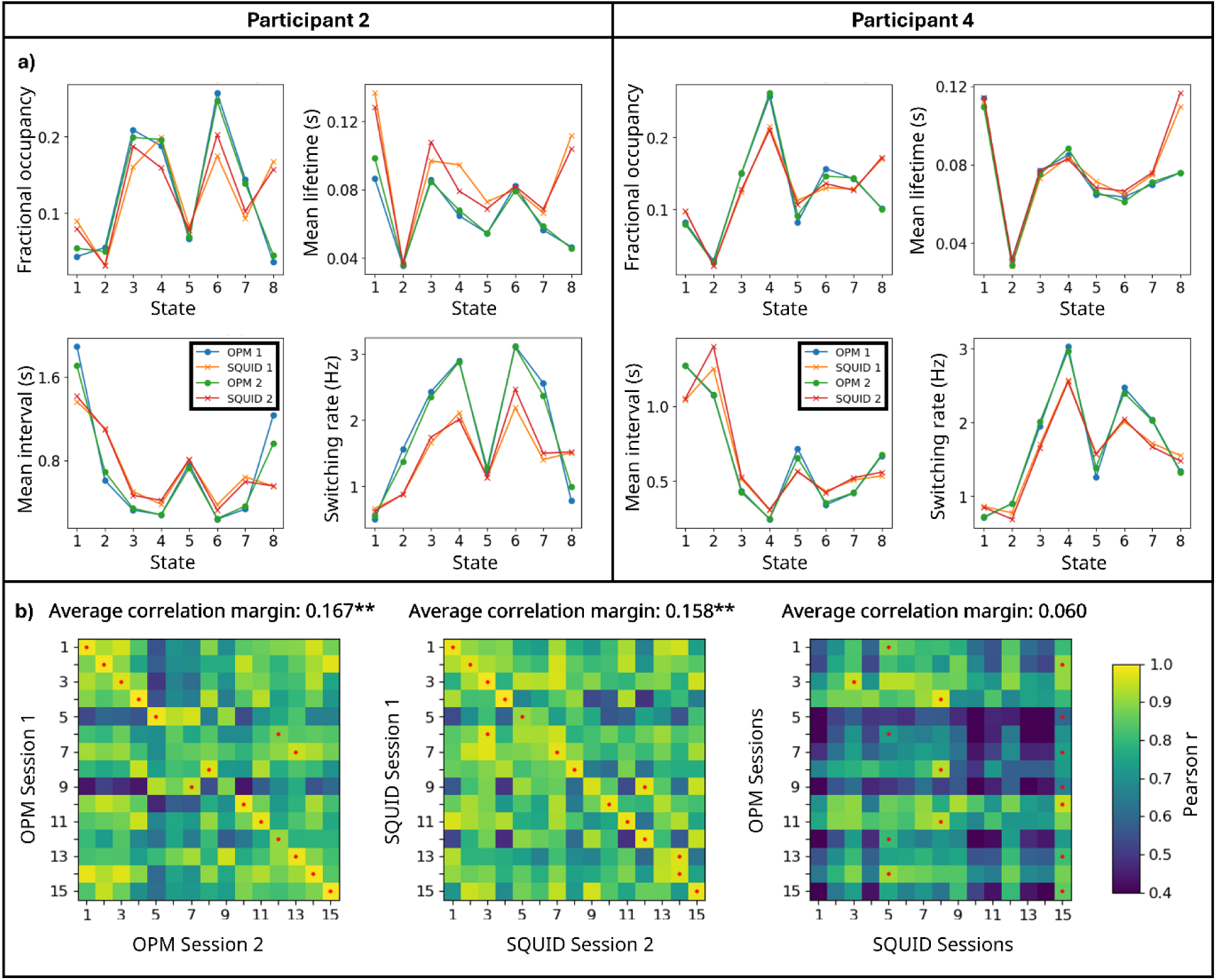
Fingerprinting using state summary statistics: a) State summary statistics (fractional occupancy, mean lifetime, mean interval time, and switching rate) shown for all states and for two representative participants. Data shown for all 4 recording sessions. b) Fingerprinting matrices comparing state summary statistics; OPM recordings (left), SQUID recordings (middle), and between modality recordings (right). Average correlation margin values are shown above each matrix; ** indicates significance following correction for multiple comparisons using Bonferroni correction.

**Figure 7:**
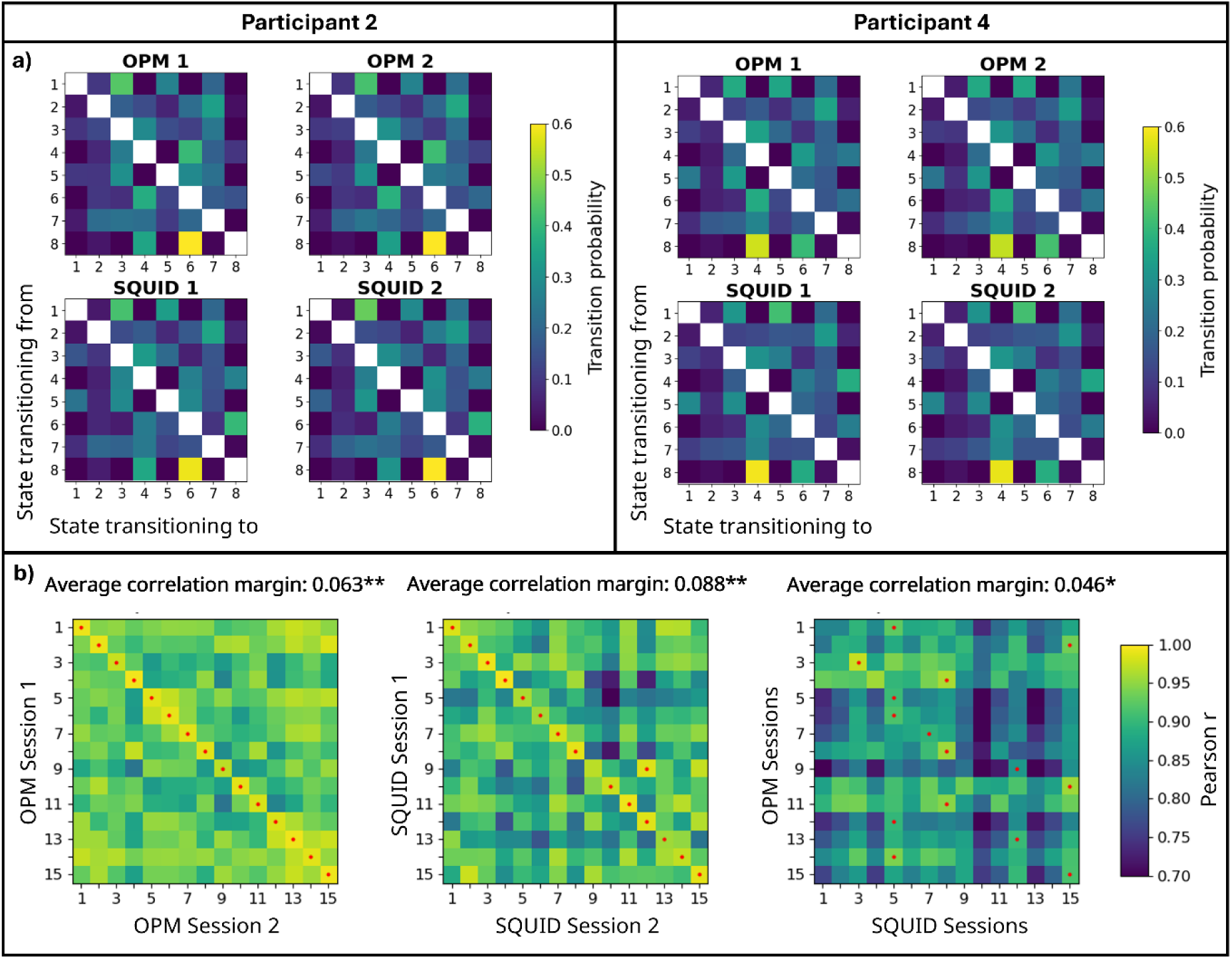
Fingerprinting using state transition matrices: a) State transition matrixes shown for two representative participants. Data shown for all 4 recording sessions. b) Fingerprinting matrices comparing state transition matrices; OPM recordings (left), SQUID recordings (middle), and between modality recordings (right). Average correlation margin values are shown above each matrix; ** indicates significance following correction for multiple comparisons using Bonferroni correction. * Indicates significance but doesn’t survive multiple comparison correction.

Interestingly, for both the summary statistics and the state transition probabilities, although there is excellent consistency between values derived within a subject and modality, there is significant variation not only between subjects, but also between modalities. For example, in Figure 6 we see that in both participants presented, the fractional occupancy of state 8 is consistently higher for SQUID-based recordings than for OPM-based recordings. The same is true for the mean lifetime of state 8 (this was also observed in the data presented in Figure 2).

The fingerprinting matrices show that, for both metrics, the within modality fingerprinting works well. For the summary statistics; 12/15 participants were correctly identified in our OPM-OPM comparison and, likewise, 12/15 (albeit a different 12) were correctly identified in our SQUID-SQUID comparison. Both average correlation margins for these within-modality comparisons were statistically significant. For the state transition matrices, all 15 participants were correctly identified in our OPM-OPM comparison and 14/15 were correctly identified in our SQUID-SQUID comparison. Again, the corresponding average correlation margins were statistically significant. However, for the between modality fingerprinting, neither the summary statistics nor the state transition matrices offered a statistically significant average correlation margin metric. This was reflected by the OPM-SQUID fingerprinting matrices, with just 3/15 participants correctly identified using the summary statistics and 5/15 correctly identified for the state transition matrices. This shows that, whilst there are individual characteristics captured using both metrics by a single modality, those characteristics did not persist between modalities. This observation will be addressed further in our discussion.

## DISCUSSION

The ability to derive features in MEG data that uniquely identify individuals is a useful means to investigate variation across a population. Moreover, it offers significant potential for discovery of ‘precision’ biomarkers specific to that individual (i.e. at what point in a disorder does an individual no longer exhibit their neural fingerprint, and can this be taken as an early marker of disease) (da Silva Castanheira et al. 2024). However, fingerprinting using MEG is a relatively new field, and this is made more complex by recent advances in MEG hardware (the introduction of OPMs) and mathematical modelling (the introduction of techniques to capture dynamic data features). Here, we have shown that individual information persists not only across SQUID and OPM-based MEG platforms, but also across static and dynamic representations of MEG data.

The application of the CHMM to both OPM and SQUID data warrants discussion. The CHMM (Gohil C et al. 2026) was trained entirely on a normative dataset captured using SQUID technology. Specifically, the normative data were collected using a MEG device comprising 204 radially-oriented planar gradiometers (17 mm baseline) and 102 radially-oriented magnetometers. This architecture is substantially different to our OPM system, which captures neuromagnetic fields using 64 radially-oriented magnetometer based channels and 128 tangentially-oriented magnetometer-based channels. It is also substantially different to our SQUID architecture, which comprised 275 radially-oriented gradiometers (5 cm baseline). Despite these marked differences, when inferring the canonical states on our data, we found that the VFEs for both the SQUID and OPM data fell within the range observed in the training data. Interestingly, despite different SQUID-based architectures, the VFE values for our SQUID data were almost identical to the training data, demonstrating that the CHMM model generalises across SQUID platforms. The VFE values for the OPM data were significantly higher than for both the training data and our SQUID data (indicating a worse fit). This is not surprising because 1) OPMs are positioned closer to the scalp, giving greater (relative) sensitivity to shallower sources. 2) OPMs have improved spatial resolution, potentially meaning an altered leakage profile between parcels) and 3) OPMs have distinct noise profiles and are susceptible to different sources of interference. These differences combined likely contribute to the significant difference in VFE values. Nevertheless, the difference in fit between OPMs and SQUIDs, whilst significant, remains marginal and the VFE values for OPM data still fell within the range of the training set. This implies that the CHMM model also generalises across the two MEG platforms. Nevertheless, the finding also highlights the need for future work to incorporate OPM MEG data into training datasets.

Our study showed that fingerprinting is possible both within and between modalities using static data features, based upon regional (parcel-level) PSDs. This is in strong agreement with previous studies (e.g. (da Silva Castanheira et al. 2021)) using conventional MEG, and supports previous fingerprinting results (Tanner et al. 2025) using the same OPM-MEG and SQUID-MEG data. However, our previous results using these data (Tanner et al. 2025) were based on images showing the spatial signature of beta modulation in sensorimotor cortex, and time frequency spectrograms of oscillatory modulation extracted from the motor regions. (I.e. fingerprinting was based entirely on sensorimotor responses to the task). Distinctly, the analyses here show that individuals features are measurable in PSD metrics taken from the whole brain (not just the sensorimotor cortices). The cross-platform nature of this finding is also important. It is easy to conceive how within modality fingerprinting could be affected by artifacts. (For example, in a conventional scanner, two people could have identical brain activity, but if one always sat with their head leaning on the left side of the scanner helmet, fingerprinting would be possible (due to signal to noise ratio differences) even though brain activity is the same). However, the fact that the unique features of individuals persist across two different MEG platforms suggests that fingerprinting has a neural (rather than artefactual) origin.

The application of the CHMM imposes a structure onto a set of MEG data. Specifically, for each time point in a new dataset, the technique determines which of the 8 canonical states (derived from normative data) is most likely to be occurring. Given this, it’s tempting to speculate that the CHMM could remove information that is specific to individuals, and only capture features of a MEG dataset that are common across a large cohort of individuals (i.e. making fingerprinting less likely). However, our results show that this is not the case. We found that both the trial-averaged state activation time courses and the state-specific PSDs independently supported successful fingerprinting both within and between MEG platforms. These measures describe when states occur, how states respond to stimulation, and the spectral composition of states. Successful fingerprinting using these metrics therefore demonstrates that individuals possess characteristic state descriptions (the spectral make up). Further, the way the canonical states dynamically respond to stimulation is unique to individuals. Overall, this implies that neural identity is not only encoded in oscillatory properties but also in the way that canonical networks – characterised by transient and punctate activations - are constructed and respond to stimulation. This finding extends previous work showing that HMM-states capture meaningful aspects of cognition and behaviour, by demonstrating that they also contain stable subject-specific information.

Although cross platform fingerprinting was possible, the OPM- and SQUID-derived datasets differ in several respects, including sensor-brain distance, spatial sensitivities, and noise characteristics. It is reasonable to assume that these differences inherently alter the absolute detectability of specific states. Indeed, this proved to be the case, with lower VFE (as noted above) a generally increased occurrence of state 8 (and a consequent drop in occurrence of the other states) for the SQUID data compared to the OPM data. These platform-induced differences were found to be larger than differences between subjects. These modality-induced differences are a likely explanation for the stark divergence in cross-modal fingerprinting performance that was observed. In particular, while state probability time courses successfully preserved individual identity across modalities, state summary statistics (and transition probabilities) did not. State summary statistics, such as fractional occupancy, rely on absolute magnitudes and are therefore highly vulnerable to the platform-specific offsets described above, which ultimately mask the individual’s neural signature. Conversely, stimulus-locked state probability time courses are largely insulated from this issue because their cross-modal similarity is evaluated using Pearson correlation. By standardizing each state’s time course independently - effectively standardising the data prior to comparison - the correlation calculation mathematically largely compensates for the absolute hardware offset. This isolates and preserves the temporal shape of the individual’s unique neural response, allowing the dynamic fingerprint to survive across disparate MEG platforms. The conclusion is therefore that neural identity is better characterised by the relative structure of state dynamics rather than absolute measures. However, future work should seek to understand how OPM-MEG and SQUID-MEG differ in the absolute likelihood of occurrence (i.e. the FO’s) of the canonical states.

There are several limitations of this study that should be considered. Firstly, the study examined a single experimental paradigm with a relatively short time between scans (all 4 scans were carried out on the same day). It therefore remains unclear whether the same pattern of results would be observed using other tasks (or in the resting-state), or with longer intervals between scanning sessions. This should be the topic of future investigation. Second, whilst correlation-based fingerprinting provides an intuitive measure of identifiability, alternative machine-learning approaches to compare two or more datasets may provide a more sensitive assessment of subject-specific information contained within state dynamics. These should be considered for future studies.

Finally, the CHMM represents a conceptual shift in the way MEG data are analysed. Traditional approaches treat each participant’s data in isolation, extracting features independently from a single recording. In contrast, the CHMM leverages information from hundreds of previously acquired datasets to learn a set of canonical brain states that capture the common spatial, spectral and temporal structure of electrophysiological activity. Here, we have shown that this framework translates across platforms, both to SQUID-MEG with a different architecture, and to OPM-MEG (albeit with a marginally, but significantly, lower VFE). Further, the technique preserves information unique to individuals. As OPM technology continues to mature, this representational framework could become increasingly valuable, providing a common basis for comparing established and emerging MEG platforms. Moreover, beyond methodological applications, the ability to characterise how canonical brain states change across the lifespan or in neurological and psychiatric disorders has considerable potential to advance our understanding of brain dynamics in health and disease.

## CONCLUSION

Using both SQUID and OPM technology, we investigated whether fingerprinting was possible both within and between MEG platforms. Further, we questioned whether fingerprinting was possible using both static and dynamic representations of MEG data, the latter being based on the CHMM. Results show that, consistent with previous work, fingerprinting was successful within and between modalities using static data features (PSDs). The CHMM was applied successfully to both modalities, yielding similar trial-averaged responses. Fingerprinting was successful using several CHMM-derived dynamic data features, including state activation time courses and state power spectral densities. However, the CHMM derived state summary statistics and transition probabilities only supported within modality fingerprinting. Together, our findings suggest that subject-specific information is preserved within canonical brain states and across MEG technologies. However, the individual features captured within absolute dynamic metrics are less robust across recording modalities.

## ACKNOWLEDGEMENTS

This work was supported by the Engineering and Physical Sciences Research Council (EPSRC) (Grant number EP/Z535722/1) and the Science and Technologies Facilities Council (Grant number UKRI601). We also acknowledge funding from Innovate UK and the UK Quantum Technology Hub in Sensors Imaging and Timing (QuSIT), funded by EPSRC (EP/Z533166/1).

